# Primary cilia regulate mechanical adaptation of tendon and enthesis via hedgehog signaling

**DOI:** 10.64898/2026.04.25.720837

**Authors:** E. Zhang, A. Feng, L. Liang, A. Moonilall, F. Fang

## Abstract

**Background:** Physical loading mediates postnatal growth, homeostasis, and healing of the tendon and its attachment to bone, which is critical for rotator cuff functional integrity. Our prior studies have highlighted the mechano-sensing role of primary cilia; However, the mechanisms through which cilia convert mechanical stimuli into structural functional adaptation under altered loading conditions remain unanswered.

**Methods:** Publicly available scRNA-seq datasets of mechanically loaded human patellar tendon cells were re-analyzed to identify cilia-related transcriptional changes. Tendon-specific cilia knockout mice (ScxCre;Ift88^fl/fl^) and wild-type controls (Ift88^fl/fl^) underwent mechanical unloading induced by botulinum toxin A injection, followed by micro–computed tomography, biomechanical testing, histology, qPCR, and immunohistochemistry to evaluate structural, mechanical, and Hedgehog (Hh) signaling responses. Primary tendon fibroblasts from wild-type and cilia-deletion mice were treated with Hh agonist or antagonist to assess Hh signaling responsiveness in vitro. Student’s t-test for two groups and two-way ANOVA for two groups with two treatments were performed for our statistical analysis.

**Results:** Here, we find that mechanical force causes changes in cilia- and hedgehog (Hh)-related gene expression in human tendon fibroblasts. Cilia ablation in the enthesis blunts force-driven remodeling of tissue structure and mechanical strength. Cilia deletion also leads to impaired Hh signaling in tendon cells and decreased responsiveness to activation and inactivation of hedgehog signaling.

**Conclusions:** Our results demonstrate loading-regulated ciliary Hh signaling during postnatal growth of the tendon and enthesis and provide proof-of-concept for developing new cilia-targeted mechanical and biological therapies for enthesis repair.

## 1. Introduction

The effects of mechanical loading on musculoskeletal tissue health and function are generally considered a double-edged sword. Physiological levels of mechanical loading are required for tissue development, homeostasis, and remodeling after injury, but pathological levels of loading promote tissue microdamage, ruptures, and loss of function in rotator cuff tissues (1). Although rotator cuff tears are among the most common upper extremity disorders in the United States, it is still challenging to prevent rotator cuff pathogenesis, achieve effective tissue regeneration, and restore full, pain-free motion (2). Although physical therapy is the predominant treatment modality, it has mixed effects on rotator cuff healing, depending on loading intensity, frequency, tear size, and patient age (1, 3, 4). At the root of this is a lack of systematic understanding of the cellular mechanisms underlying how cells sense and convert mechanical signaling, adjust their function accordingly, and subsequently alter macro-level tissue structure and composition.

An increasing body of work in the musculoskeletal field suggests that the primary cilium, an organelle present in most mammalian cells, functions as a mechanical and chemical sensor of the extracellular environment. As mechanosensors, primary cilia adjust their length and prevalence in response to changes in force (5, 6). Therefore, in vivo cilia deletion dampens or abolishes loading-stimulated tissue formation and healing (5, 7-10). Consistently, our previous studies have shown that the tendon-bone attachment (enthesis), whose injury or degeneration mainly contributes to rotator cuff tear, modulates cilia incidence in response to different loading scenarios (6). Genetic disruption of ciliary protein expression causes abnormal enthesis structure and decreased mechanical function. However, mechanistic insights into the essential role of primary cilia in tendon and enthesis mechano-transduction in vivo remain limited.

The ciliary membrane concentrates numerous signaling proteins and lipids that modulate responses to cellular and environmental cues. One of the most well-explored and enthesis-related ciliary signaling pathways is hedgehog (Hh) signaling. When Hh ligand binds to its receptor Ptch1, Ptch1 leaves the cilium, allowing Smo to enter the cilium and ultimately activates the GLI family member transcription factors like GLI1 and GLI2 (11). During enthesis growth, Hh-responsive cells with stemness features form graded fibrocartilage in the enthesis and Smo-deficient mice display defected layout of extracellular matrix (6, 12). During enthesis repair, Hh activation via pharmaceutical administration increases fibrocartilage mineralization and further improves biomechanical strength (13). All this evidence shows the potential role of cilia-centered mechanotransduction and transport of Hh signaling during enthesis growth and repair. However, little is known about whether cilia are required for enthesis mechano-responses and Hh-stimulated cell behaviors, which could guide the development of cilia-oriented treatment with the combined effects of both physical and biologic therapies.

In this work, we sought to determine how primary cilia mediate mechanotransduction in tendon and enthesis cells and whether they are required for regulating Hh signaling in response to mechanical loading. We hypothesize that primary cilia function as essential mechanosensors that couple mechanical stimuli to Hh signaling pathway, thereby regulating mechanical adaptation of the enthesis. We demonstrate that mechanical stimulation is associated with changes in cilia- and Hh- related gene expression in human tendon cells. Genetic ablation of primary cilia in mouse tendon and enthesis cells led to reduced Hh expression and attenuated enthesis accommodation in response to loading deprivation. Taken together, these findings indicate that primary cilia contribute to the regulation of mechanosensitive and Hh- related signaling pathways in tendon and enthesis.

## 2. Materials and methods

### 2.1 Animals

Experimental protocols involving animals were reviewed and approved by the Mount Sinai Institutional Animal Care and Use Committee. The mice were housed on a 12h light/dark cycle and fed with food and water ad libitum. To delete cilia specifically in the tendon and tendon enthesis, ScxCre transgenic mice originally provided by the Schweitzer lab from Oregon Health and Science University (14) were crossed with Ift88^fl/fl^ mice obtained from Jackson Laboratory (022409). ScxCre;Ift88^fl/fl^ mice show evidence of discernible polycystic kidney disease at the age of 8 weeks and these cilia-ablated mice have no significant difference in tendon morphology and mechanical strength before 8 weeks (6). To avoid systematic effects caused by kidney disease at the adolescent stage, ScxCre; Ift88^fl/fl^ (cKO^Scx^) mice were unloaded from postnatal day 1 (P1) to week 6. Ift88^fl/fl^ mice were used as wild-type controls (WT^Scx^). Both male and female animals were evenly and randomly selected by blinded researchers to be assigned into different treated groups, including unloading, different outcome measurements, and in vitro examination. For each experiment, the figure legends include the number of mice analyzed and the biological replicates. For mouse shoulder loading deprivation *in vivo* experiment, 9 mice were used in both the WT^Scx^ and cKO^Scx^ groups (n = 9 per group), for a total of 18 mice. For the mouse shoulder enthesis in vitro experiments, 5 mice were used per group (n = 5 per group), with a total of 10 mice. For mouse tail tendon in vitro experiments, 3 mice were used per group, with a total of 12 mice. Sample sizes were determined based on previous studies using similar experimental designs and outcome measurements. Exclusion criteria were established a priori. Animals were excluded if they showed signs of illness unrelated to the experimental procedure, if the treatment delivery was unsuccessful, or if tissue samples were damaged during processing and could not be reliably analyzed. Data points were also excluded if technical failures occurred during mechanical testing or imaging that prevented accurate measurement.

### 2.2 scRNA-sequencing analysis of human tendon cells

scRNA-seq datasets of three human healthy patellar tendons with and without in vitro loading/stretching were accessed as processed cellRanger files from public repositories (GEO accession GEO: GSE150482). The control and loaded datasets were integrated and reanalyzed following the Seurat single-cell analysis pipeline(15). Briefly, low quality cells with <500 genes and mitochondrial contamination >20% were excluded from each dataset, followed by data normalization and scaling. Due to unavailable prior knowledge about the specificity of primary cilia on cell types and avoidance of biased analysis, Graph-based clustering and UMAP dimensionality reduction were performed to take all cells as one cluster and the control and loaded can be separated. Differentiated expressed genes (DEGs) were identified using the Seurat FindMarkers function with default parameters. Enriched biological pathways were detected based on these DEGs using Enrichr. Top enriched biological processes and cellular components related to cilia biogenesis and function were reported. To account for potential technical variation between the control and loaded samples, we employed well-established bioinformatics normalization pipelines, including Seurat integration and Harmony (16, 17). By using the Local Inverse Simpson’s Index (LISI) as a key evaluation metric for cell-type conservation (16), we identified Seurat reciprocal principal component analysis (RPCA) as the optimal approach to identify anchors and align shared cell populations across different experimental batches (17). Furthermore, data scaling was conducted using log-normalized and centered counts to minimize batch effects while preserving robust biological shifts in the transcriptomic state.

### 2.3 Unloading models

Botulinum toxin A (BtxA)-mediated unloading was performed as previously described, which has been validated to result in decreased muscle volume, decreased bone volume in humerus head, decreased deposition of fibrocartilage tissue at the enthesis, and impaired enthesis mechanical strength (6, 18-20). Specifically, one shoulders of cKOScx, WT^Scx^ mice received 0.2 U BtxA in saline from the date of birth until 6 weeks. The contralateral shoulders were used as control and received the same amount of saline. The tendon-enthesis-humerus head construct was harvested at the end of 6 weeks and analyzed by the approaches outlined below (21, 22)

### 2.4 Tendon fibroblast isolation and treatment

Tail tendon from WT^Scx^ and cKO^Scx^ mice were isolated and chopped in pieces. Tissues were placed in a 15 ml conical tube with 5 ml sterile phosphate-buffered saline containing 2 mg/ml collagenase type II (Worthington) and 1X penicillin/streptomycin (Gibco) for two hours. Afterwards, the digested tissues were filtered via 100 μm strainer and cultured until confluency in alpha-MEM with 10% fetal bovine serum (Benchmark) and 1X penicillin/streptomycin in a humidified incubator of 37°C and 5% CO2.

For Hh agonist and antagonist treatment, WT^Scx^ and cKO^Scx^ cells were plated and cultured in 12-well plates until 60% confluency, starved overnight, and treated with 100 mM Hh agonist Hh-Ag1.5 for 8 hours and 100 uM Hh antagonist cyclopamine for 6 hours. mRNA was isolated and analyzed by qPCR.

### 2.5 Biomechanical testing

The supraspinatus tendon-bone complex from WT^Scx^ and cKO^Scx^ mice were dissected and kept hydrated in PBS for mechanical testing. The humerus head was placed in a customized 3D-printed fixture and tendon was clamped in steel grips (6, 21). The tissue and grip assembly was mounted on an Electroforce testing frame (TA instruments) with a bath of PBS at 39°C. The samples were loaded between 0.05N and 0.2N for cycles, recovered for 3 minutes, and stretched to failure with 0.2%/S. The recorded force and strain data was used to determine enthesis structure and material properties.

### 2.6 Microcomputed tomography

Dissected supraspinatus tendon-to-bone construct were examined for bone morphometry as described previously (6). Microcomputed tomography (Bruker Skyscan 1272) was set with an energy of 55 kilovolt peaks, an intensity of 145 μA, and a resolution of 5 μm to scan the samples. Cross-sectional areas were measured from the constructed 3D images. The humeral head between enthesis and the growth plate was selected by a segmentation algorithm and analyzed for parameters of cortical and trabecular bone quality.

### 2.7 Histology and immunostaining

Dissected supraspinatus tendon-bone complex was fixed in 10% neutral buffered formation (Sigma), decalcified in Versenate Decal solution (StatLab) for 14 days, and embedded in optimal cutting temperature compound (6). Enthesis sections were collected using a cryostat and stained by safranin O following the standard procedure. For immunostaining, the sections were pretreated with 2 mg/ml hyaluronidase for 1 hour and incubated in 0.5% Triton X. 15% goat serum was used for section blocking for 1 hour. The primary antibodies including Ihh (Proteintech,13388-1-AP, 1:400), Shh (R&D systems, AF464-SP, 1:20), GLI1 (Abcam, ab49314, 1:500), were used to incubate the sections overnight. Fluorescence-conjugated secondary antibodies (ThermoFisher) with appropriate hosts and species reactivity were applied for 1 hour. After washing with PBS three times to remove extra primary antibodies, the sections were mounted with VECTASHIELD antifade mount medium containing DAPI (Vector Laboratories) and sealed with nail polish.

Images of Z-stacks were acquired on a Zeiss LSM 980 equipped with Airyscan 2 using a 20X or 60X objective. The researchers were blinded to tissue genotypes and groups during image collection and analysis. The images were analyzed by ImageJ software. The percentage of positive cells for each sample was calculated across 3 fields with at least 100 cells.

### 2.8 qPCR analysis

mRNA expression related to cilia assembly, hedgehog signaling, Wnt signaling, tenogenesis, and fibrocartilage formation were analyzed using qPCR. Following mice euthanasia, the supraspinatus entheses were immediately dissected under a micro-dissection microscope, snap-frozen in liquid nitrogen, and mechanically pulverized. RNA was isolated by TRIzol (Life Technologies) and cleaned using spin-columns (PureLinkd RNA mini kit, Thermo Fisher Scientific). Reverse transcription was conducted using the High-Capacity cDNA Reverse Transcription Kit (Invitrogen). The cilia-focused gene expression was evaluated using RT2 Profile PCR Assay for mouse primary cilia (Qiagen) by following the product manual. All gene expression data were normalized to the housekeeping genes, then to the expression of WT entheses, and presented as 2−ΔΔCt. For in vitro cell experiments, mRNA was isolated, cleaned, and reversely transcribed as described above. The genes were examined by SYBR Green-based approach (Invitrogen) using a Quant Studio 6 Flex (Applied Biosystems). The gene expression was firstly normalized by the housekeeping gene glyceraldehyde 3-phosphate dehydrogenase (GAPDH), followed by normalization of gene expression from the control group of tendon fibroblasts without Hh agonist or antagonist treatment.

### 2.9 Statistics

The conditions of each experiment were blinded during data collection and analysis. GraphPad Prism was used for two-sided statistical tests. Student’s t-test for two groups and two-way ANOVA for two groups with two treatments were performed for our statistical analysis. After ANOVA analysis, we adjusted p values of multiple comparisons using Sidak’s method. All data and plots are shown as mean±□standard deviation.

## 3. Results

### 3.1 Mechanical loading regulates ciliation in tendon cells

Our previous studies showed that mouse tendon and tendon enthesis cells assembled primary cilia, and cilium assembly was mediated by physical activity (6). However, it is not clear how primary cilia function as mechano-sensors in human tendon cells to adjust their composition and thereby adapt to different physical conditions. Therefore, we re-analyzed publicly available single-cell RNA sequencing (scRNA-seq) data of isolated healthy human patellar tendon cells after loading in vitro (23) (Fig. 1a). After unsupervised clustering using UMAP, most loaded cells were clustered together with an obvious separation from cells without loading (Control, Fig. 1b). Since the identities of tendon cells with primary cilia are unknown, genes that were differentially expressed in tendon cells due to loading were identified by comparing transcriptional profiles of all the cells with and without loading. We found that ARL2, UBE2L3, and AURKAIPI, which are involved in ciliary protein trafficking (24, 25) and deacetylation of ciliary axonemal microtubules (26), were downregulated in loaded cells (Fig. 1c-d). This was also true for TUBB6, IFT57, and DYNLL1, which encode proteins required to assemble cilia (27-29). Since Hh signaling, one of the most well-recognized ciliary signaling pathways (30), has been found to regulate tendon and tendon enthesis development, injury responses, and homeostasis, we also evaluated whether loading would alter expression of Hh-related transcriptional genes. Although a subpopulation of unloaded cells showed low expression of hedgehog-related genes GLI1, GLI2, and GLI3, showing cell heterogeneity, a remarkable number of unloaded cells expressed these proteins at higher expression levels, consistent with the up-regulation of ciliary genes in unloaded cells (Fig. S1A-B). Gene ontology (GO) enrichment analysis of genes upregulated in unloaded human cells further showed over-representation of genes associated with the ciliary landscape and enhanced ciliary molecular function (i.e., ubiquitin conjugated enzyme activity, cyclase activator activity) and cellular component (i.e., organelle inner membrane, Fig. 1e). Collectively, our scRNA-seq data revealed a significant downregulation of ciliary structural and functional genes in mechanically loaded human tendon cells. While these transcriptomic changes suggest that primary cilia respond to mechanical force, scRNA-seq data remain inherently observational. Inspired by the observation, we would conduct further animal work to functionally validate the role of cilia in mechanical sensing.

**Fig. 1.**
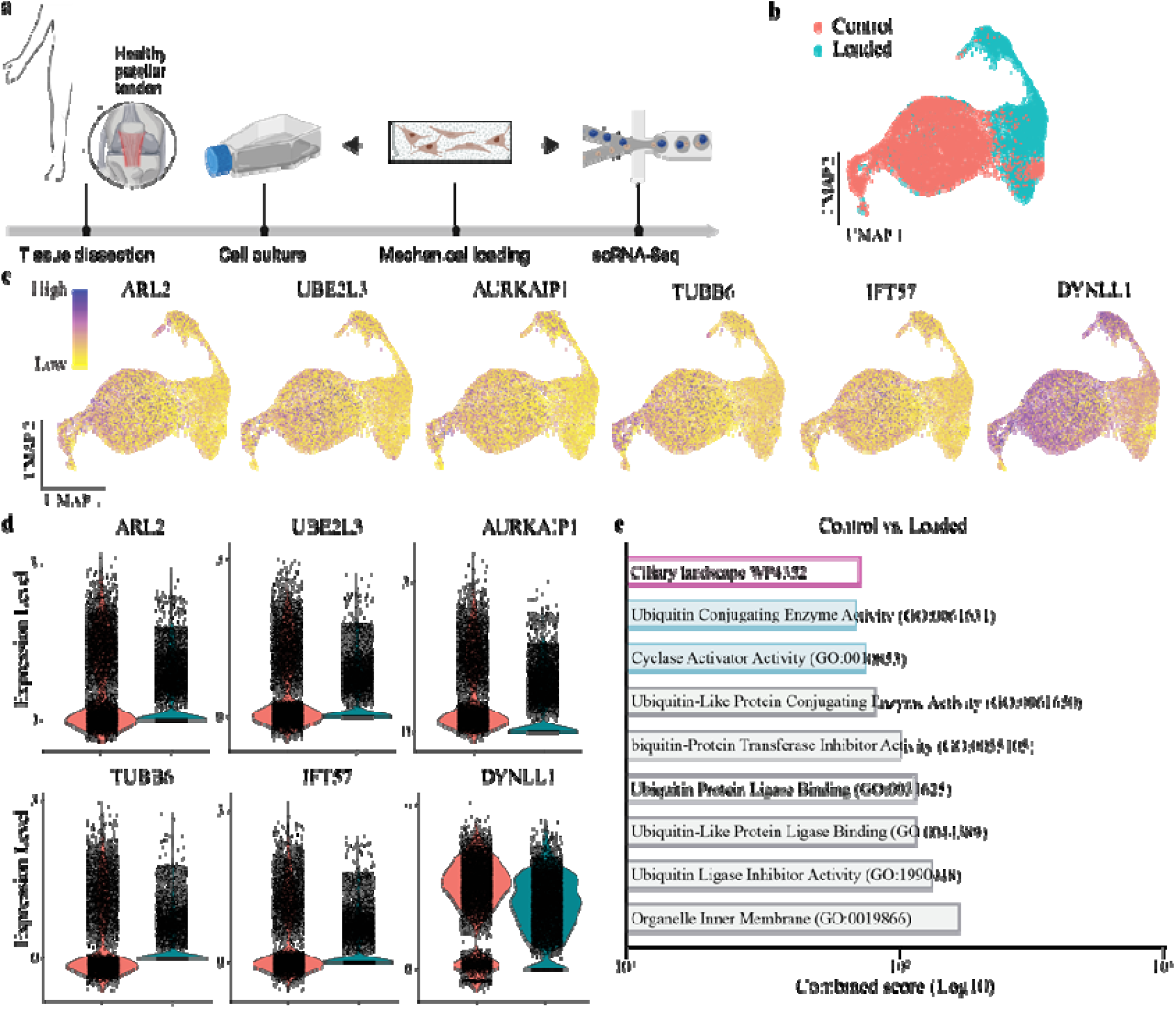
Mechanical loading regulates genes related to ciliation in human tendon. a scRNA-seq analysis of human tendon cells after harvest, in vitro expansion, and mechanical loading. b UMAP plot of control (red) and loaded (blue) tendon cells. Visualization of genes involved in cilia function (i.e., ARL2, UBE2L3, AURKAIPI), and ciliary structural genes (i.e., TUBB6, IFT57, DYNLL1) of control and loaded tendon cells by feature (c) and violin (d) plots. e Over-represented biological processes and cellular components of control tendon cells compared to the ones after loaded, identified by GO enrichment analysis.

### 3.2 Cilia disruption attenuates tendon-to-bone attachment mechanical adaptation

IFT88 is essential for cilium assembly, and we previously created a conditional Ift88 knockout mouse model wherein primary cilia were ablated in tendon and tendon-enthesis cells using ScxCre (6), which directs Cre expression specifically in these cell types. Cilium formation was significantly reduced in enthesis cells, and altered enthesis composition and mechanical strength were observed when compared to their wildtype (WT^Scx^) controls (6). The dramatically altered skeletal phenotype and the inability of knockout mice to perform treadmill-running tasks prevented our use of loading models in our conditional knock-out mice. Therefore, we focused on the effects of unloading on wildtype (WT^Scx^) and cilia knockout mice (cKO^Scx^, Fig. 2a), where unloading was realized by injection of botulinum toxin A in the shoulder. First, the quality of bone between the head of the humerus and growth plate was examined. For both WTScx and cilia-depleted mice, unloading significantly decreased cortical volume, trabecular bone volume, and trabecular bone density (Fig. 2b), confirming effective unloading and preserved bone remodeling, as primary cilia remain intact in bone tissue. Cortical bone density was not altered after unloading in WTScx or cilia-depleted mice. WTScx mice also had similar cortical bone volume, trabecular bone volume, and trabecular bone density compared to knockout mice in both control (cage activity) and unloading conditions. Since cilia were only specifically removed in the tendon and tendon enthesis, similar responses to unloading were expected in the bone of WTScx and knockout mice. However, these observations did not hold for the supraspinatus enthesis. WTScx mice exhibited a significant decrease in cross-sectional area, maximum force, and work to yield after unloading (Fig. 2c), (p<0.05), but unloading did not result in the corresponding differences in these mechanical properties in knockout mice. Mechanical parameters, including Young’s modulus, stiffness, and ultimate stiffness of WTScx and knockout mice were not significantlyaltered after unloading (Fig. 2c and S2a). Consistently, WTScx mice had enthesis mechanical properties comparable to knockout mice in control and unloading conditions. This suggests that cilia depletion from embryonic to early postnatal stages may not directly affect tendon enthesis growth (6). Additionally, cilia-mediated mechanotransduction potentially regulates enthesis formation and maturation. Morphologically, unloading caused less fibrocartilage (in red) deposited at the enthesis of WTScx mice, which was not observed in knockout mice (Fig. S2b). Unnoticeable fibrocartilage in red was found from the cKOScx supraspinatus entheses in both the control and unloading scenarios. WTScx mice also had more round chondrocytes with relatively bigger lacuna in cage activity, compared to after unloading, but cKOScx mice from both the control and unloading groups had elongated chondrocytes with neglectable lacuna. These results demonstrate that in the absence of primary cilia, the tendon enthesis is unable to alter its structure, composition, or mechanical properties to accommodate different loading states.

**Fig. 2.**
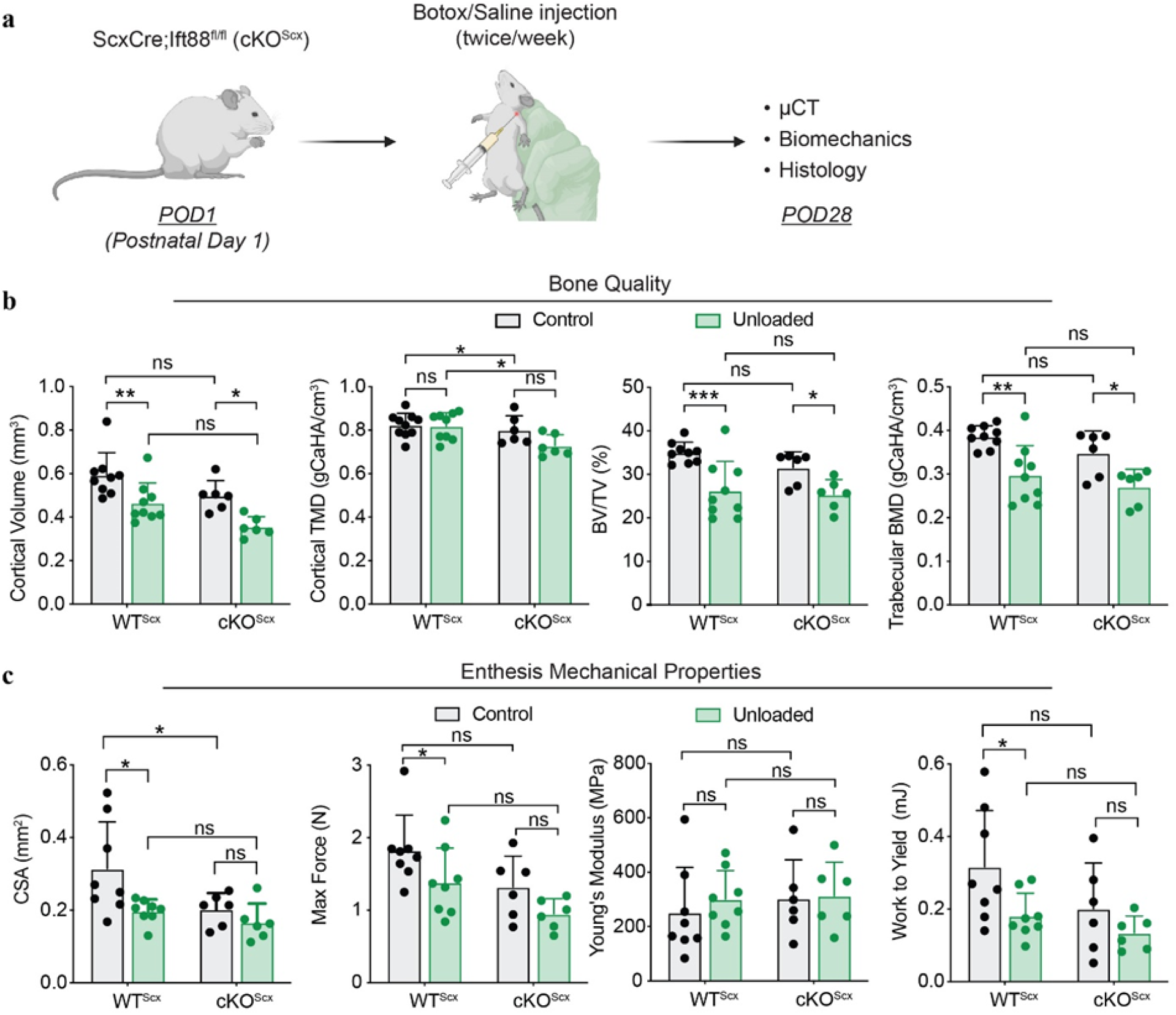
Cilia ablation dampens tendon enthesis mechanical responses. a Generation of supraspinatus tendon enthesis of cilia-deleted transgenic mice with loading deprivation by injecting botulinum toxin A (BtxA). ScxCre;Ift88fl/fl, cKOScx. Created with BioRender.com. b Humeral head bone quality from WTScx (Ift88fl/fl) and cKOScx. TMD, tissue mineral density; BV/TV, bone volume fraction; BMD, bone mineral density. c Tendon enthesis mechanical properties of WTScx (Ift88fl/fl) and cKOScx measured by mechanical testing. CSA, cross-section area; Max Force, maximum force; during mechanical testing until enthesis failure. N=9 mice/group. Data are shown as mean ± SD. Two-way ANOVA analysis with multiple comparisons using Sidak multiple comparisons test is performed. *, p<0.05; **, p<0.01; ***, p<0.001. ns, no significant difference.

### 3.3 Loss of cilia exhibits abnormal Hh signaling in vivo

Primary cilia have been reported to be the signaling center for multiple signaling pathways, including, but not limited to, TGFβ signaling, Hh signaling, and Wnt signaling (11). To identify cilia-mediated cellular mechanisms relevant to this tissue, we performed qPCR and immunohistochemistry on tendon enthesis from cilia knockout mice (Fig. 3a). Interestingly, Genes of Hh ligands, such as Ihh and Shh, which are prevalent during skeletal development, were more highly expressed in the cKOScx enthesis than the WTScx enthesis, especially with Shh showing a significant difference. Nevertheless, GLI1 as a key transcription factor for Hh signaling activation, was significantly less expressed in the cKOScx group compared to the control (Fig. 3b). Since Hh receptors and effectors are located on the cilium surface, we conclude that the elevated production of Hh ligands represents a compensatory response to diminished Hh signaling caused by reduced ciliation. However, this compensatory mechanism was unable to rescue the defects in Hh signaling, reflected by decreased percentage of GLI1-positive cells. Similarly, cilia ablation also led to upregulation of the Wnt signaling ligand, Wnt9b, but no difference was observed in Wnt downstream genes. Given that GLI1-expressing cells have been found to function as progenitor or stem cells by populating the whole region of tendon enthesis and differentiating into, for example, enthesoblasts, mineralizing chondrocytes, and unmineralizing chondrocytes to construct enthesis (21). Therefore, we focused on cilia-regulated Hh signaling in the enthesis for this study. We confirmed that the ratios of cells expressing Ihh or Shh protein in cKOScx mice did not differ from those in WTScx mice, but less cells exhibited GLI1 expression in the tendon enthesis of cKOScx mice (Fig. 3c). These findings indicate that the role of primary cilia in enthesis growth operates primarily through ciliary dependent Hh signaling.

**Fig. 3.**
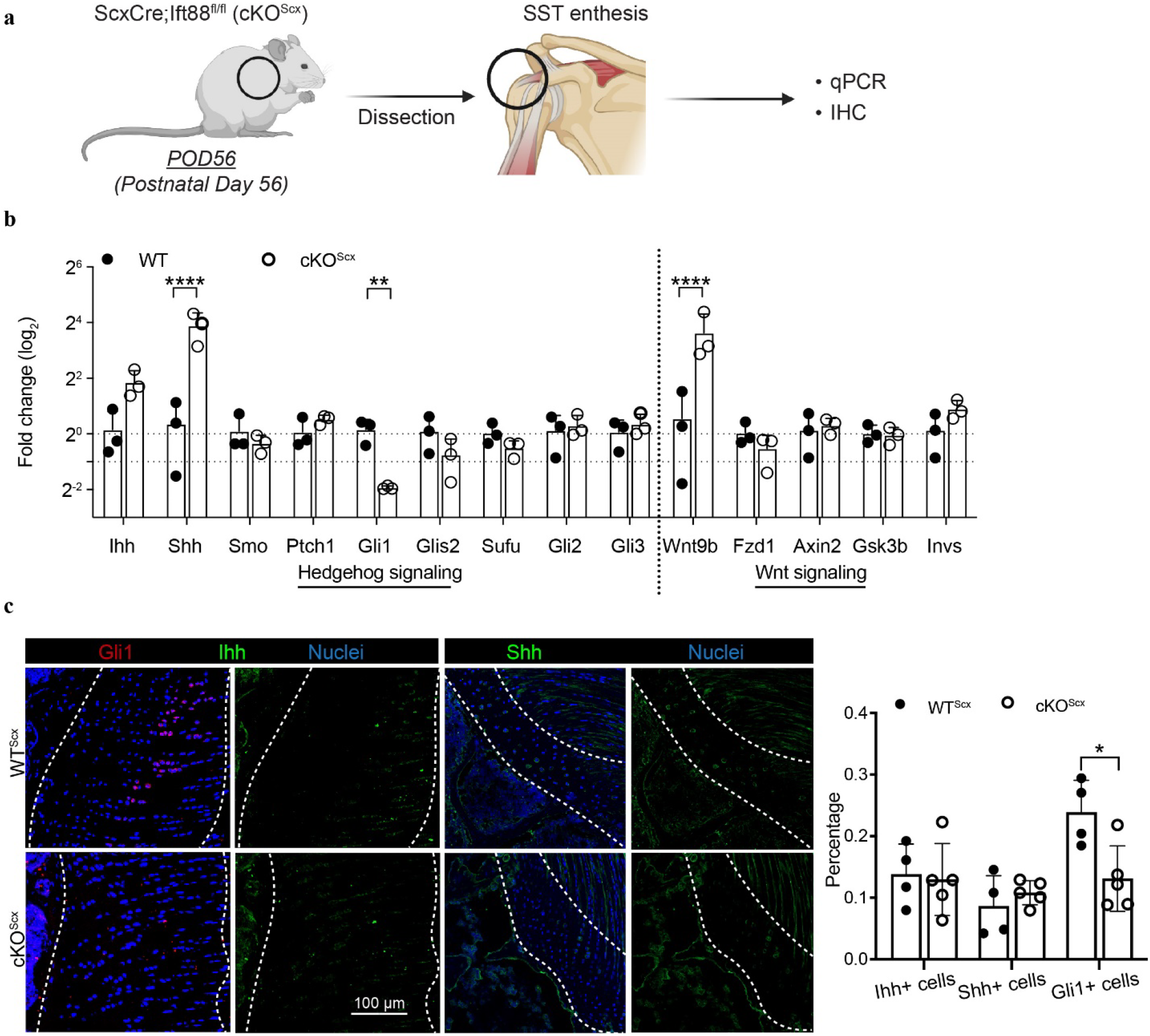
Primary cilia are required for Hh signaling in tendon and tendon enthesis. a WTScx and cKOScx supraspinatus tendon enthesis dissected and used for evaluation of gene and protein expression. Created with BioRender.com. b q-PCR analysis of Hh- and Wnt-related genes in tendon enthesis of WTScx and cKOScx mice at 8 weeks. c IHC staining (left) of Ihh, Shh, and GLI1 in eight-week-old WTScx and cKOScx mice and quantification of the percentage of cells (right) expressing the corresponding Hh-related proteins. n=3 mice/group in (b), and n=5 mice/group in (c). Data are shown as mean ± SD. Unpaired Two-sided student’s t test for data analysis in (b) and (c). *p< 0.05, **p<0.01, and ****p< 0.0001. IHC, immunohistochemistry; POD, postnatal day.

### 3.4 Cilia deletion attenuates Hh signaling responsiveness in tendon cells in vitro

To further evaluate whether altered expression of Hh ligands by cilia loss or less presented Smoothened on cilia membrane contributes to ciliary Hh signaling, we treated tail isolated tendon fibroblasts from cKOScx and WTScx groups with Hh agonist Hh-Ag1.5 and Hh antagonist cyclopamine (31, 32), both of which target Smoothened (Smo, Fig. 4a). Tail tendon cells, as valuable sources which have been widely used to characterize tendon fibroblast behavior, were selected as an alternative to supraspinatus tendon cells, due to the lower cell density and more heterogenous cell population (21). To eliminate baseline differences in gene expression between cKOScx and WTScx groups (Fig. S3a), we normalized the expression of each gene after Hh agonist or antagonist treatment by the expression level of the same gene without treatment for both cKOScx and WTScx groups. As expected, cKOScx cells expressed less Hh related genes Patch1 and GLI1, compared to WTScx cells (Fig. 4b). However, genes related to cilium assembly, tenogenesis, and fibrocartilage formation, were similar between cKOScx and WTScx groups after Hh activation. Ciliation and GLI1 expression were not differentially affected in cKOScx and WTScx cells (Fig. S3b, c). When inhibited with Smoothed antagonist, cKOScx cells were more resistant, compared to WTScx cells, as reflected by higher GLI1 expression (Fig. 4c). All other genes related to cilia assembly, tenogenesis, and fibrocartilage formation were not differentially expressed between cKOScx and WTScx cells, except for significantly less expression of tenomodulin in cKOScx cells. Consistently, at the protein level, cKOScx and WTScx groups exhibited similar ciliation, but GLI1 protein expression was also elevated in cKO group (Fig. S3b, c). These results may imply that highly differentiated tendon fibroblasts are potentially stable and resistant to in vitro Hh activation and inactivation, which might limit transcriptional changes in their gene expression during tenogenesis and fibrocartilage formation. Our observations that cilia-depleted cells display dampened responses to Hh activation and inactivation via Smoothened regulation, indicate that Hh signaling is directly controlled by ciliation or cilia lengthening in this tissue.

**Fig. 4.**
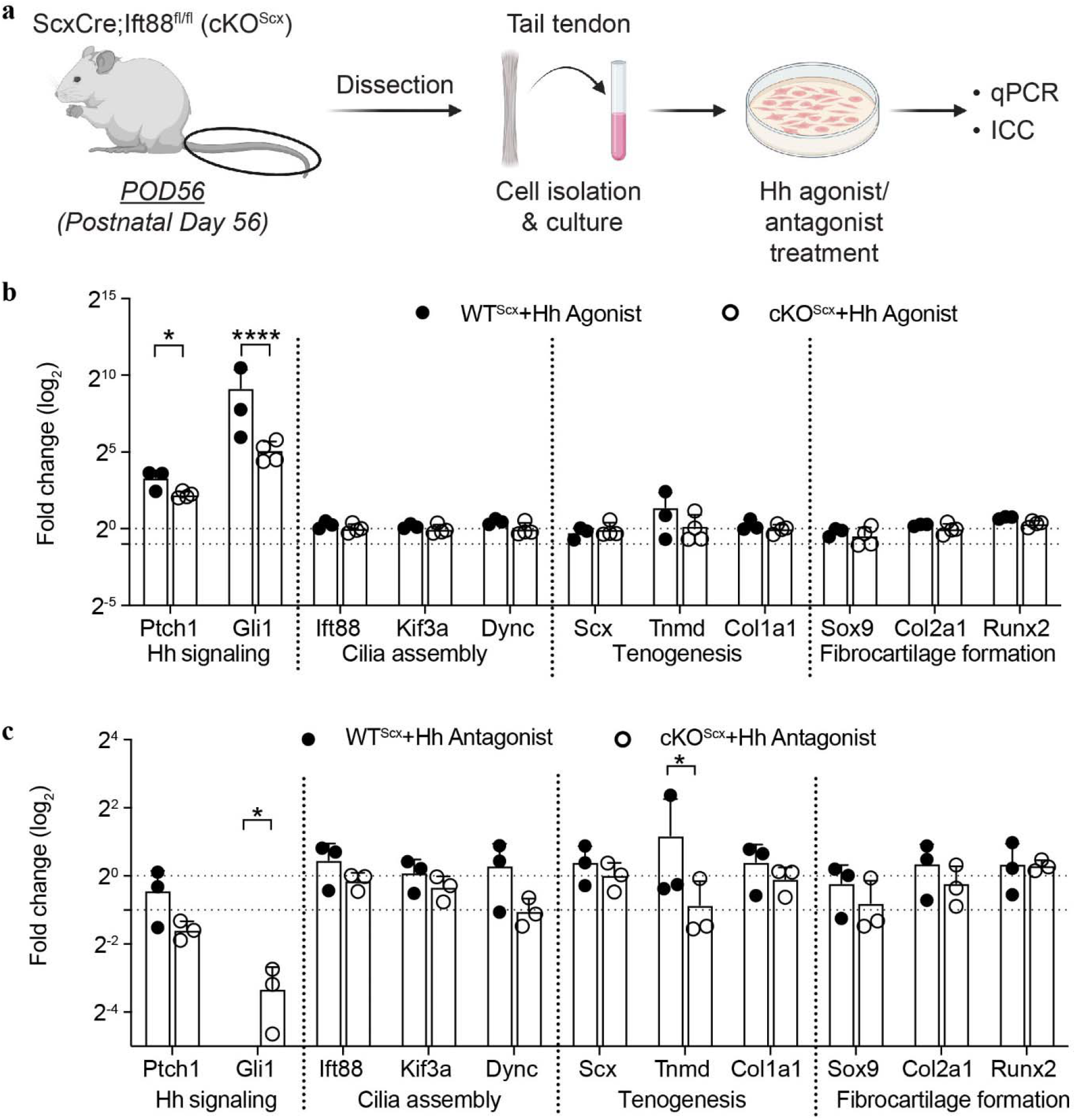
a Tail tendon from these mice used for cell isolation and in vitro evaluation. Created with BioRender.com. b, c q-PCR analysis of eight-week-old tail tendon cells from WTScx and cKOScx mice after treatment with 100 nM Hh agonist Hh-Ag1.5 or 10 µM Hh antagonist cyclopamine for 8 hours. The gene expression of WTScx and cKOScx cells in (b) and (c) after agonist or antagonist treatment are normalized by the corresponding gene expression without treatment to eliminate the effects caused by cilia deletion. n=3 mice/group in (b), and (c). Data are shown as mean ± SD. paired t-test for analysis in (b) and (c). *p< 0.05, **p<0.01, and ****p< 0.0001. POD, postnatal day.

## 4. Discussion

Mechanical regulation of tendon and tendon enthesis has been examined in terms of cell morphology, constituent, structure, and both mechanical and material properties. The identification and characterization of mechanosensation and transduction at the cellular level would help harness the power of physical therapy to promote tissue healing and regeneration. Our work indicates the regulation of the loading-cilia-Hh signaling axis during postnatal growth of the tendon enthesis: mechanical loading affects ciliogenesis, and primary cilia, in turn, control tissue mechanical adaptation and Hh signaling activation.

Cilium biogenesis has been found to be species- and tissue-dependent. Most recent cilia research focused on animal tissues and organs. While only a limited number of reports demonstrate the presence and ratio of primary cilia in human orthopedic tissues and their function in vitro and ex vivo (33). This study conducted an unbiased analysis of human scRNA-seq data and identified downregulation of several cilia-related genes after loading, which is consistent with our previous results showing decreased cilia-related gene expression and cilia presence in mouse cells. Supporting this, in vitro cyclical loading was also found to cause primary cilia disassembly in human tendon cells (34). Additionally, in vitro loading also downregulated Hh-related genes in tendon cells, consistent with findings from animal work (6). The evolutionary conservation of primary cilia function across diverse species indicates the high potential of translating and applying experimental findings from animal models to develop clinically relevant cilia-targeted treatments for musculoskeletal tissue regeneration.

Our work demonstrates that primary cilia control Hh signaling during postnatal enthesis growth. Ciliary gene Ift88 deletion caused an increase of Hh ligand Shh gene expression but decrease of Hh transcription factor GLI1. It is possible that increased ligand production represents an attempt to compensate for downregulation of Hh signaling in entheses. This compensatory effect was insufficient to recover normal Hh signaling, as the proportion of Shh- and Ihh-positive cells remain largely unchanged following cilia deletion, while the percentage of the GLI1-positive cell was reduced. Besides Hh signaling, cilia were found to mediate Wnt signaling, shown by increased Wnt ligand after cilia loss. The role of Wnt signaling has not been well addressed in tendon and enthesis. Limited data have proposed that Wnt signaling would maintain tendon stem cells, repress expression of tendon markers, and promote tendon healing (35-37). Therefore, the effects of cilia deletion on enthesis structure and function are potentially driven by a combination of well-studied (i.e., Hh signaling) and under-studied (i.e., Wnt signaling) cellular pathways, highlighting the importance of a detailed understanding of cilia-centered physiology.

Hh signaling has been identified as a key therapeutic target to promote rotator cuff healing. The development of a library of small molecules such as Hh agonists will further accelerate their repurposing in musculoskeletal disorders. Of note, ciliogenesis must be well tuned while manipulating Hh signaling by treatments with Hh agonists or antagonists. Our observation that cellular responses to Hh activation or inactivation were diminished significantly with dysfunctional cilia demonstrated the importance of the interaction and realignment of ciliary signals and cilia function for therapeutic treatment. Furthermore, different cell types and tissues demonstrated distinct responses to Hh stimulation and cilia-mediated regulation, which illustrates that the development of therapeutic drug options centered around cilia and Hh signaling should be cell type-dependent. We reported that tendon tail fibroblasts, with or without primary cilia, had similar transcriptional expression of genes associated with tenogenesis and fibrocartilage formation after successful activation of Hh signaling pathway. One possible explanation is that tendon tail fibroblasts represent a relatively mature and stable cell population that is less responsive to environmental intervention.

This study is not without limitations. First, tendons from different types and species were used across different experimental models, which may introduce variability related to differences in cellular composition and mechanical environment. In addition, this study employed a constitutive ScxCre;Ift88fl/fl model with analyses restricted to early postnatal stages. Future studies will use inducible, tissue-specific deletion strategies to examine ciliary function in mature tendon and enthesis.

## 5. Conclusions

While cilia have been found to regulate TGF-β, Hh, Notch, and Wnt signaling across various tissues, whether and how these different pathways are coordinated by cilia to mediate tendon or enthesis function is unclear. Additionally, given that activation of mechanical loading or Hh signaling has been proved to increase enthesis healing, targeting cilia for enthesis repair might have the potential to activate both biomechanical and Hh signaling for enhanced tissue repair. Our study identifies a mechano-reulatory axis linking mechanical loading, primary cilia, and Hh signaling in tendon enthesis growth and adaptation. Supporting this, our analysis of human scRNA-seq data indicates that mechanical stimulation is associated with changes in tendon cell ciliogenesis. Furthermore, our in vitro and in vivo results suggest that tissue adaptation to mechanical loading and Hh signaling activation at the enthesis is modulated by primary cilia. Therefore, our study would provide the insight into cilia as a hub for synchronizing biomechanical and hedgehog cues and cilia as a better therapeutical target for rotator cuff treatment.

## Supplementary Material

**Supplementary Fig. 1.**
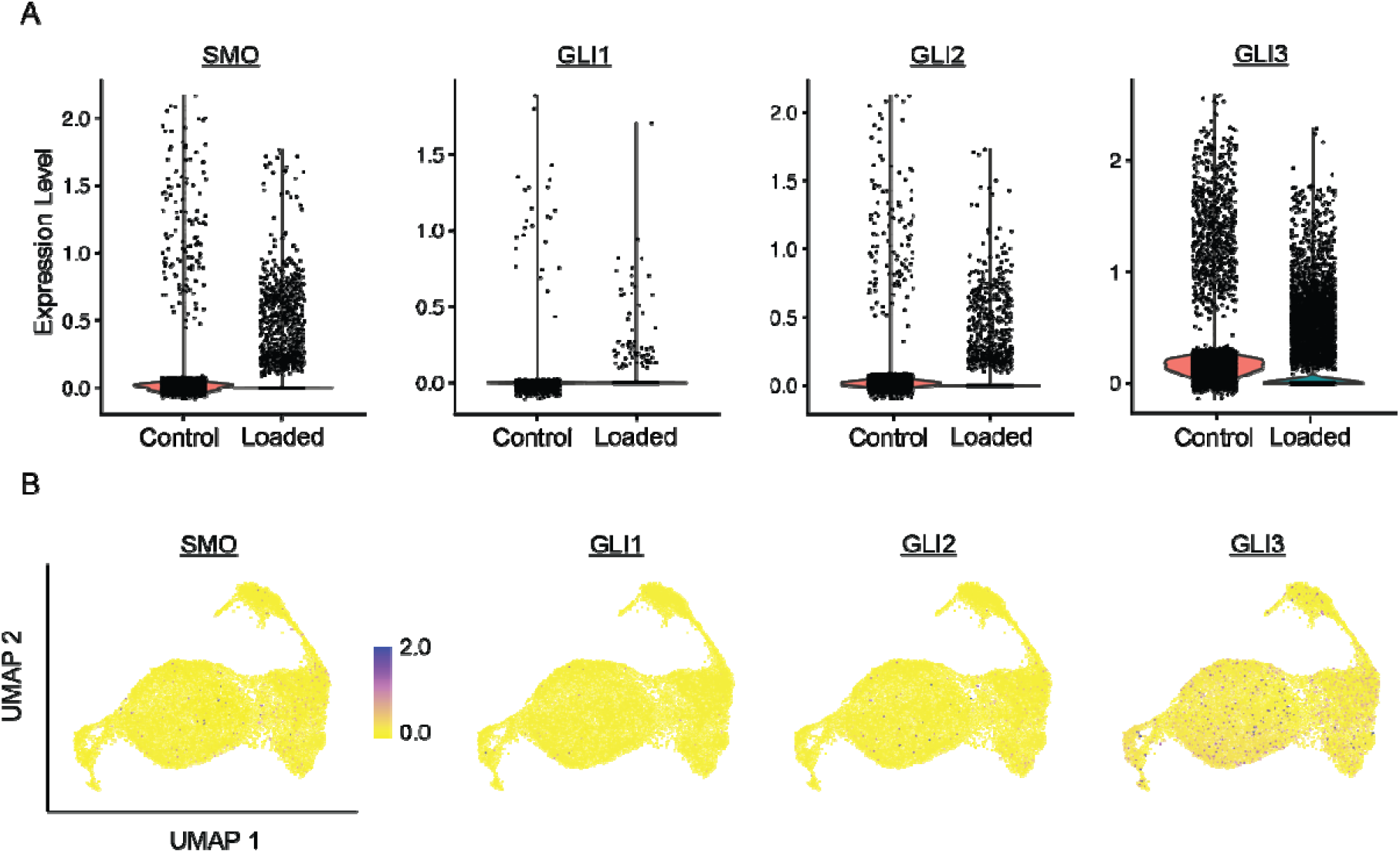
Expression of Hh-related genes (i.e., SMO, GLI1, GLI2, GLI3) in human control and loaded samples shown by (A) violin plots and (B) feature plots.

**Supplementary Fig. 2.**
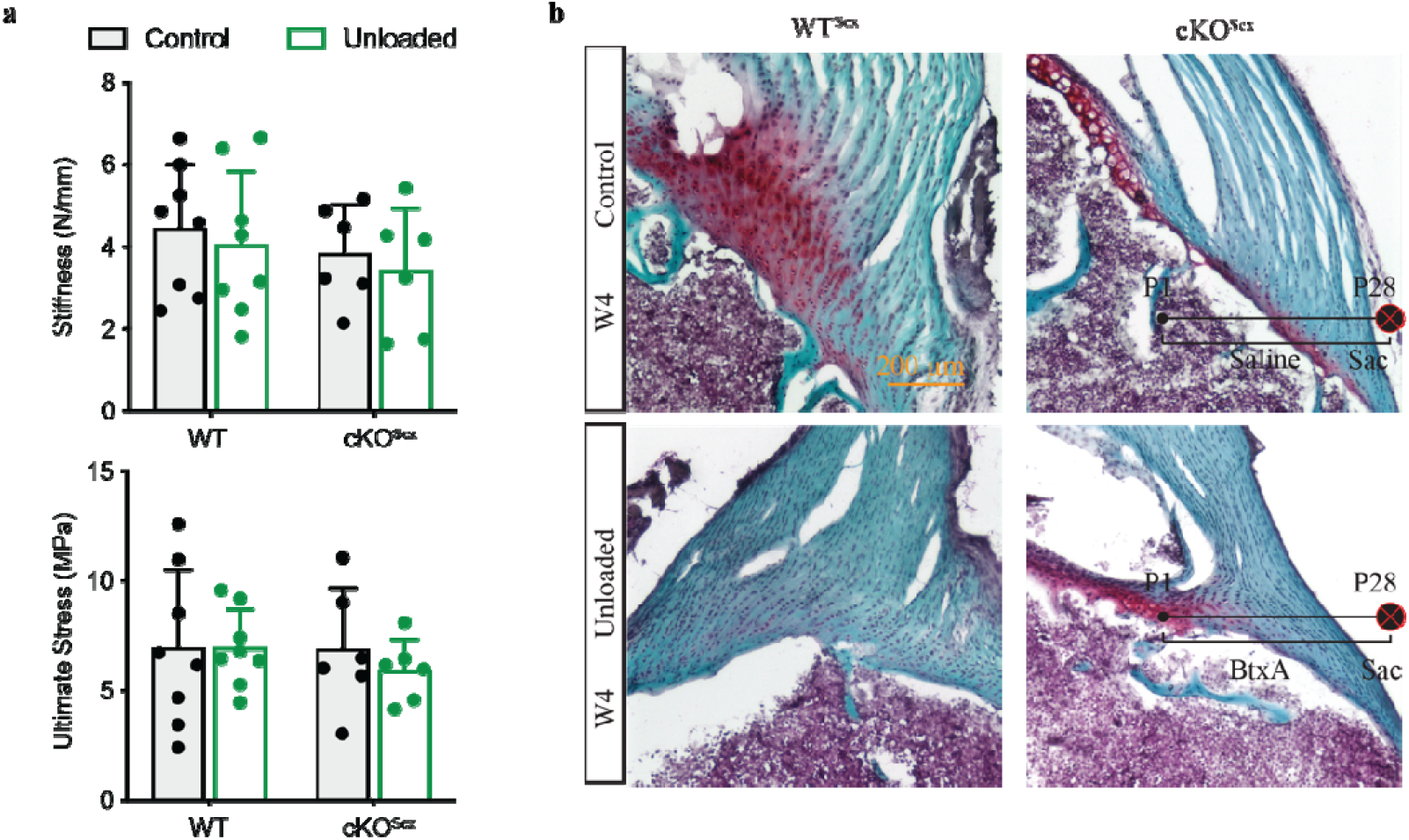
Cilia deficiency blocks loading-induced tissue remodeling. a Tendon enthesis stiffness and stress from WTScx (Ift88fl/fl) and cKOScx (ScxCre;Ift88fl/fl) mice measured by mechanical testing. b Safranin O staining of the tendon enthesis showing decreased fibrocartilage region (red) of WTScx mice after unloading compared to control, but cKOScx mice showing negligible structure difference of the tendon enthesis after unloading. N=6-8 mice/group in (a) and (b). Data are shown as mean ± SD. Two-way ANOVA analysis with multiple comparisons using Sidak multiple comparisons test was performed.

**Supplementary Fig. 3.**
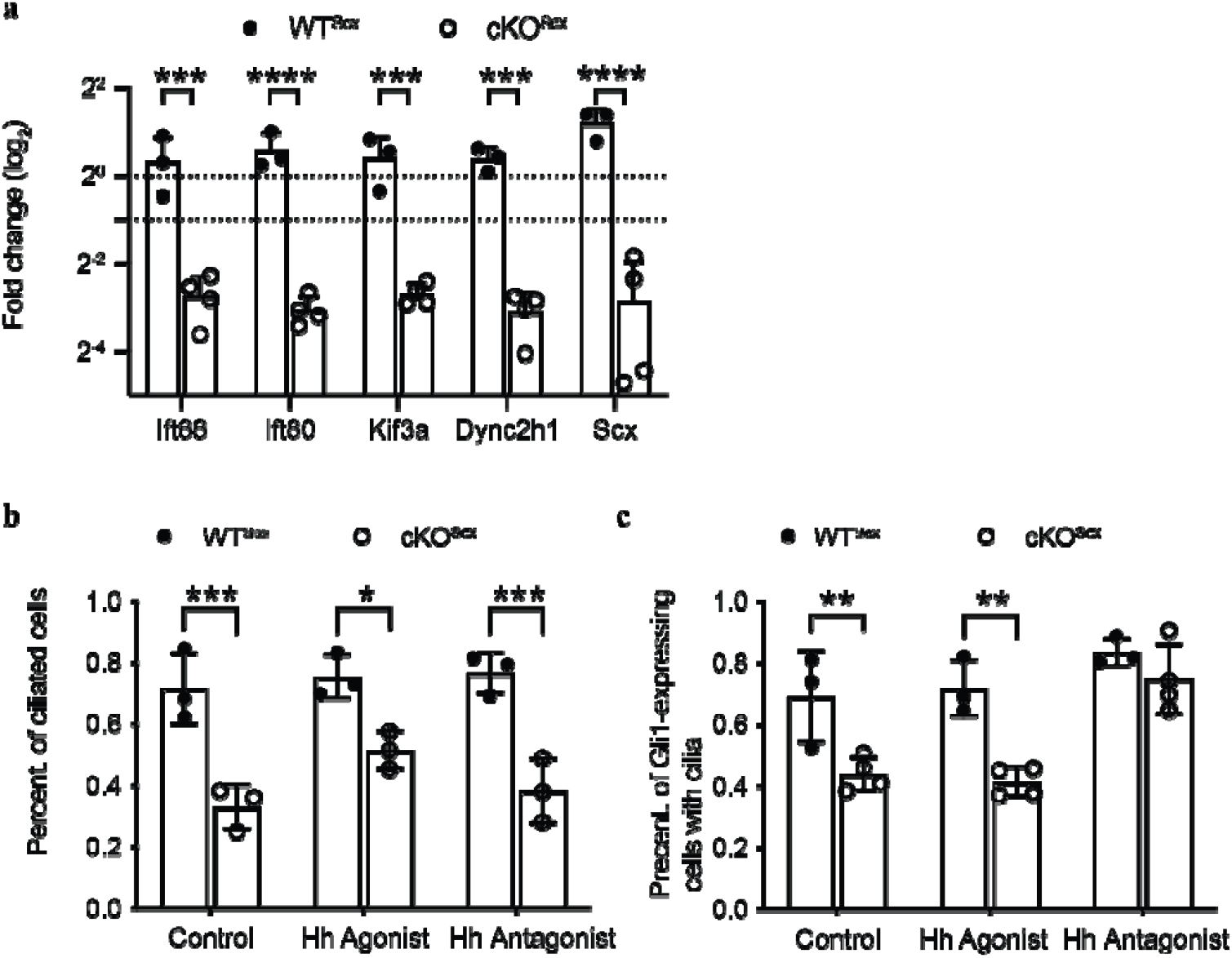
Ciliation is independent on Hh signaling in tail tendon cells in vitro. a Tail tendon cells from cKOScx (ScxCre;Ift88fl/fl) mouse models have significantly decreased cilia-related (i.e., Ift88, Ift80, Kif3a, Dync2h1) and tenogenic (i.e., Scx) genes, compared to WTScx (Ift88fl/fl) mice, demonstrating knockout efficiency in cilia. b, c Immunostaining analysis of tail tendon cells treated with Hh agnonist Hh-Ag1.5 and Hh antagonist cyclopamine shows both the percentages (percent.) of ciliated cells (normalized by the total cell number) and GLI1-expressing cells with cilia (normalized by the cell number of GLI1-expressing cells) decrease in cKOScx cells, but ciliation is muted from Hh activation or inactivation. This analysis is conducted on fluorescently staining tail tendon cells after Hh agonist or antagonist treatment using acetylated tubulin (targeting cilia) and GLI1 antibodies. n=3-4 mice/group. Data are shown as mean ± SD Unpaired Two-sided student’s t test for data analysis is performed in (a) and two-way ANOVA analysis with multiple comparisons using Sidak multiple comparisons test is conducted in (b) and (c). *, p<0.05; **, p<0.01; ***, p<0.001.

## Availability of Data and Materials

All data are available in the main text or the supplementary materials. Source data are provided within this paper.

## Author Contributions

Conceptualization: F.F.; Methodology: E.Z., A.F., L.L. and A.M.; Investigation: E.Z., A.F., L.L. and A.M.; Visualization: E.Z. and F.F.; Supervision: F.F.; Writing (original draft): E.Z. and F.F.; Writing (review & editing): E.Z., A.F., L.L., A.M. and F.F..

## Ethics Approval and Consent to Participate

Animal care and handling were conducted according to NIH animal care guidelines at the Icahn School of Medicine at Mount Sinai. The protocols were reviewed and approved by the Institute Animal Care and Use Committee (D16-00069).

## Acknowledgment

We thank the Microscopy and Advanced Bioimaging Core Facility at the Icahn School of Medicine at Mount Sinai for technical support.

